# Paraventricular Thalamus Hyperactivity Mediates Stress-Induced Sensitization of Unlearned Fear but Not Stress-Enhanced Fear Learning (SEFL)

**DOI:** 10.1101/2025.05.30.657116

**Authors:** Kenji J. Nishimura, Denisse Paredes, Nathaniel A. Nocera, Dhruv Aggarwal, Michael R. Drew

## Abstract

Exposure to stress can cause long-lasting enhancement of fear and other defensive responses that extend beyond the cues or contexts associated with the original traumatic event. These nonassociative consequences of stress, referred to as fear sensitization, are thought to underlie some symptoms of trauma-related disorders. Fear sensitization has been predominately studied using the Stress-Enhanced Fear Learning (SEFL) paradigm, which models the stress-induced amplification of fear learning. Less is known about the mechanisms through which unlearned fear responses are sensitized by stress. Here, we investigated the neural mechanisms for sensitization of unlearned fear responses using a paradigm we termed Stress-Enhanced Fear Responding (SEFR). In this model, mice exposed to a single session of footshock stress exhibit enhanced freezing to a novel tone stimulus. To investigate brain regions that might mediate SEFR, we first used c-Fos mapping to identify neural activity changes associated with stress-induced enhancement of unlearned fear. Our c-Fos screen identified the posterior paraventricular thalamus (pPVT) as a region that was persistently hyperactive after footshock stress and whose activity correlated with behavioral expression of SEFR. Using fiber photometry, we observed that SEFR, but not SEFL, was associated with increased activity in the pPVT. Next, we found that chemogenetic inhibition of the pPVT blocked both the induction of SEFR during stress and its later expression, while artificial stimulation of pPVT in stress-naive mice was sufficient to recapitulate SEFR. Interestingly, pPVT inhibition or stimulation did not affect acquisition or expression of SEFL. In conclusion, our results indicate that sensitization of fear learning (SEFL) and sensitization of unlearned fear (SEFR) have distinct neural mechanisms. Our results identify pPVT hyperactivity as a mechanism for stress-induced sensitization of unlearned fear and highlight pPVT as a potential target for treating arousal and reactivity symptoms of trauma- and stressor-related disorders.

## Introduction

Exposure to severe stress can lead to the development of psychiatric conditions such as post-traumatic stress disorder (PTSD). While PTSD encompasses a range of symptoms, alterations in fear learning are widely regarded as a core feature underlying the disorder (***Lissek and Meurs, 2015***). Accordingly, Pavlovian fear conditioning, a form of associative learning, has been the dominant framework for studying how fear is triggered by cues or contexts associated with a prior stressful experience (***Bienvenu et al., 2021***). However, stress also engages a second mechanism often overlooked in Pavlovian procedures: fear sensitization, a nonassociative form of learning. Fear sensitization is characterized by a widespread potentiation of fear and defensive responses that extends beyond the initial stressful experience. Therefore, after stress exposure, innocuous stimuli may elicit fear even in contexts quite different from the original stressor. Nonassociative learning processes, such as sensitization, are highly conserved across species, from organisms with primitive nervous systems to humans (***Harris, 1943***). Sensitization is also likely to underlie a subset of PTSD symptoms (***Stam, 2007b***; ***Charney et al., 1993***; ***McFarlane, 2010***). Core symptoms—including heightened arousal and reactivity, hypervigilance, and exaggerated startle reactions—closely resemble fear sensitization. These symptoms can manifest independently of cues or contexts related to the traumatic event and may therefore be resistant to conventional therapeutics, such as exposure therapy, which primarily target associative fear processes. Despite its clinical relevance, the neural basis of fear sensitization in the mammalian nervous system remains largely unknown and underexplored.

Most research on fear sensitization has focused on a phenomenon called stress-enhanced fear learning (SEFL). Developed by Fanselow and colleagues (e.g., (***Rau et al., 2005***)), this model demonstrates that a prior history of inescapable footshock stress potentiates subsequent fear conditioning to a novel context. This phenomenon is thought to reflect stress enhancement of fear acquisition rather than fear expression. For instance, ***Rau et al. (2005***) showed that footshock stress failed to enhance the expression of conditioned fear that was acquired before and tested after exposure to footshock stress. Brain regions implicated in SEFL include the amygdala (***Perusini et al., 2016***; ***Pennington et al., 2024***; ***Daws et al., 2020***; ***Sillivan et al., 2017***), hippocampus (***Hersman et al., 2019***), and prefrontal cortex (***Pennington et al., 2017***), with the basolateral amygdala (BLA) playing a critical role. For instance, disrupting neuronal activity or protein synthesis in the BLA blocks stress-induced enhancement of fear learning (***Perusini et al., 2016***; ***Pennington et al., 2024***). In line with this, suppression of SEFL-associated microRNAs in the BLA, such as miR-135b-5p, attenuates stress effects on fear learning (***Daws et al., 2020***).

In addition to enhancing associative fear learning, stress can sensitize unlearned or unconditioned fear responses, a phenomenon we term stress-enhanced fear responding (SEFR). For example, prior footshock stress potentiates freezing evoked by a novel, innocuous tone stimulus (***Hassien et al., 2020***; ***Kamprath and Wotjak, 2004***; ***Siegmund and Wotjak, 2007a***,b). This stress-induced sensitization of unlearned fear persists even after animals undergo fear extinction in the stress context, indicating a nonassociative process that is independent of the fear memory of the initial stressful event (***Hassien et al., 2020***; ***Rau et al., 2005***; ***Siegmund and Wotjak, 2007a***). Clinically, SEFR may model the exaggerated threat sensitivity associated with PTSD (***Stam, 2007a***). However, the neural mechanisms of SEFR are poorly understood, and it is unclear whether SEFL and SEFR reflect distinct processes or share overlapping neural mechanisms.

Here, we investigated the neural mechanisms of SEFR and asked whether these mechanisms also mediate SEFL. Using c-Fos mapping and fiber photometry, we identified the posterior paraventricular thalamus (pPVT) as a region that is hyperactive in stressed animals during SEFR expression. Using chemogenetic manipulations, we show that pPVT hyperactivity during and after stress is essential for SEFR but not SEFL. We conclude that sensitization of fear learning and sensitization of unlearned fear are dissociable at the behavioral and neural levels. Additionally, we identify pPVT hyperactivity as a mechanism for stress-induced sensitization of unlearned fear and as a potential contributor to arousal and reactivity symptoms in trauma- and stressor-related disorders.

## Results

### Exposure to footshock stress induces stress-enhanced fear responding (SEFR) and stress-enhanced fear learning (SEFL)

Our laboratory previously established a mouse behavioral protocol for within-subjects assessment of SEFR and SEFL (Fig. 1A) (***Hassien et al., 2020***), based on the work of ***Rau et al. (2005***) and ***Siegmund and Wotjak (2007a***). We selected mice from the 129S6/SvEvTac strain because our previous work demonstrated that 129s6/SvEvTac mice exhibit widespread alteration in fear behaviors following exposure to footshock stress (***Hassien et al., 2020***). It should be noted that others have reported deficits in fear extinction in this substrain (***Temme et al., 2014***); however, we have previously demonstrated successful extinction learning in 129S6/SvEvTac mice under our experimental conditions (***Hassien et al., 2020***). Furthermore, ***Temme et al. (2014***) reported that 129S6 mice, in contrast to the 129S1 substrain, show reduced fear generalization and robust fear discrimination. Mice were assigned to either a Stress group, which received a single session of four 1-mA foot-shocks, or a NoStress group, which underwent context exposure for an equivalent duration without footshock (Fig. 1B). Subsequently, all mice were subjected to a series of behavioral tests designed to evaluate associative fear memory of the initial stress context, SEFR, and SEFL. The tests were conducted in order of increasing severity to minimize hysteresis.

**Figure 1.**
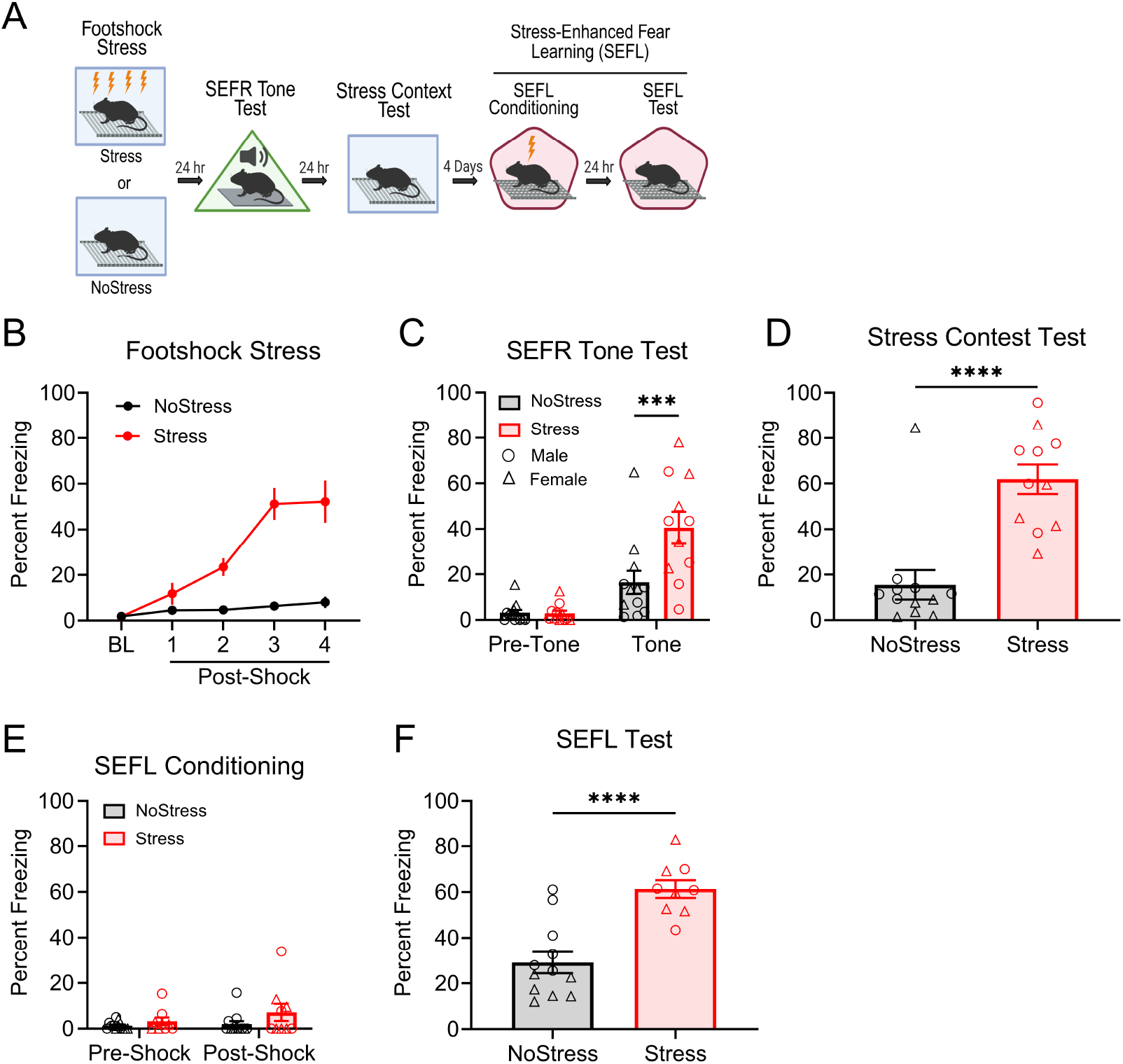
Footshock stress induces associative fear, SEFR, and SEFL. (A) Mice received a single session of footshock stress (Stress group) or were exposed to the same context and received no footshocks (NoStress group). Beginning 24 hours after stress, mice underwent the SEFR tone test, fear retrieval in the stress context, and the two-day SEFL procedure. (B) In the stress session, freezing behavior increased as a function of the 4 shock presentations. (C) In the SEFR tone test, freezing behavior was evaluated before (Pre-Tone) and during the tone and inter-tone periods (Tone). All mice exhibited low levels of pre-tone freezing. However, during the tone period, stressed mice displayed increased freezing relative to the NoStress group. (D) Stressed animals displayed robust context-elicited freezing behavior to the stress chamber. (E) During the SEFL conditioning session, all mice displayed little to no freezing prior to the onset of footshock. (F) In the SEFL test session, stressed mice exhibited increased context-elicited freezing relative to the NoStress group. Data presented as mean +/-SEM. N=23: NoStress group (6 males, 6 females), Stress group (6 males, 5 females). * *p* < 0.05, ** *p* < 0.01, *** *p* < 0.001.

To test associative fear memory, mice were re-exposed to the chamber in which the footshock stress occurred. Mice in the Stress group exhibited significantly elevated freezing levels compared to those in the NoStress group (*t*_21_ = 5.078, *p* <.0001), indicating robust associative fear memory of the stress context (Fig. 1D). To assess sensitization of unlearned fear, we employed a behavioral assay referred to as the SEFR tone test. Mice were placed in a novel chamber and exposed to three 30-sec auditory tones following a 3-min baseline period (Fig. 1C). Unlike Pavlovian cued fear conditioning, tone-elicited fear in this task reflects an “unlearned” or “unconditioned” fear response, as mice had no prior experience with the tone or the context. During the pre-tone baseline period, we observed low levels of freezing in all groups. However, during the tone and inter-tone periods (“tone” in figures), the Stress group exhibited significantly higher freezing relative to the NoStress group (*p* <.01). A two-way repeated measures ANOVA revealed a significant Group x Tone Period interaction (*F*_1,21_ = 9.870, *p* =.0049). Finally, we assessed sensitization of fear learning using the two-day SEFL procedure. On day 1 (SEFL conditioning), mice were placed in another novel context and received a single 0.75-mA footshock. All groups displayed low levels of freezing before shock delivery (Fig. 1E). On day 2 (SEFL test), mice were returned to the same context of the previous day for assessment of context-elicited fear. The Stress group exhibited significantly greater freezing than the NoStress group (*t*_19_ = 5.050, *p* <.0001), indicating enhanced fear learning (Fig. 1F).

It is important to highlight that mice displayed little to no freezing prior to the onset of the tone in the SEFR tone test (Fig. 1C “Pre-Tone”) or prior to the onset of the footshock in the SEFL conditioning session (Fig. 1E “Pre-Shock”). These observations demonstrate that mice distinguished between contexts and that subsequent freezing is these tests is not likely attributed to associative fear processes - namely, generalization of fear of the stress context. Collectively, these results demonstrate that footshock stress induced both associative fear of the stress context and fear sensitization in the forms of SEFR and SEFL. We propose that SEFR and SEFL are distinct types of fear sensitization: the sensitization of unlearned fear responses (SEFR), and the sensitization of fear learning (SEFL).

### Associative fear, SEFR, and SEFL are dissociable at the behavioral level

To determine whether associative fear, SEFR, and SEFL are dissociable at the behavioral level, we examined how the three assays respond to variations in footshock stress intensity and the extent to which freezing responses in the three assays intercorrelate. Male mice were assigned to one of four treatment groups corresponding to the amplitude of footshock delivered during the stress session: NoStress, 0.25mA-Stress, 0.5mA-Stress, and 1mA-Stress. During the footshock stress session (Fig. 2A), we detected a significant effect of Group (*F*_3,23_ = 13.04, *p* <.0001) and Group x Shock Period interaction (*F*_12,92_ = 6.384, *p* <.0001). Mice exposed to footshock showed increased levels of freezing relative to the NoStress group. In the SEFR tone test (Fig. 2B), a RM-AVOVA detected a significant Group x Tone Period interaction (*F*_3,23_ = 3.607, *p* =.0286). Pre-tone levels of freezing were low and similar across groups. During the tone period, freezing levels in the 0.5mA-Stress and 1mA-Stress groups were significantly higher than the NoStress group (all *p*′*s <*.05). Moreover, no significant differences in freezing were observed between the 0.25mA-Stress and NoStress groups. In the stress context test (Fig. 2C), results from a one-way ANOVA revealed a significant effect of Group on freezing levels (*F*_3,23_ = 30.46, *p* <.0001). Specifically, all groups that received footshock during the stress session, regardless of shock amplitude, exhibited significantly higher levels of freezing relative to the NoStress group (all *p*′*s* <.05). Lastly, in the SEFL procedure, mice were given a 1-shock conditioning session (Fig. 2D), in which significant effects were observed for Group (*F*_3,23_ = 3.413, *p* =.0344) and Group x Shock Period interaction (*F*_3,23_ = 3.439, *p* =.0336). The following day, in the SEFL test (Fig. 2E), we observed a significant main effect of Group (*F*_3,23_ = 3.391, *p* =.0351). Post hoc comparisons indicated that both the 0.5mA-Stress and 1mA-Stress groups exhibited significantly higher levels of freezing compared to the NoStress group (all *p*′*s <*.05). The pattern of results indicates that associative contextual fear had a lower threshold for induction than either form of sensitization. Lastly, we examined whether freezing behavior in stressed mice correlated across the different behavioral assays (Fig. 2F). No significant correlations were observed between associative fear of the stress context, SEFR, and SEFL, suggesting that these fear responses reflect dissociable outcomes of footshock stress.

**Figure 2.**
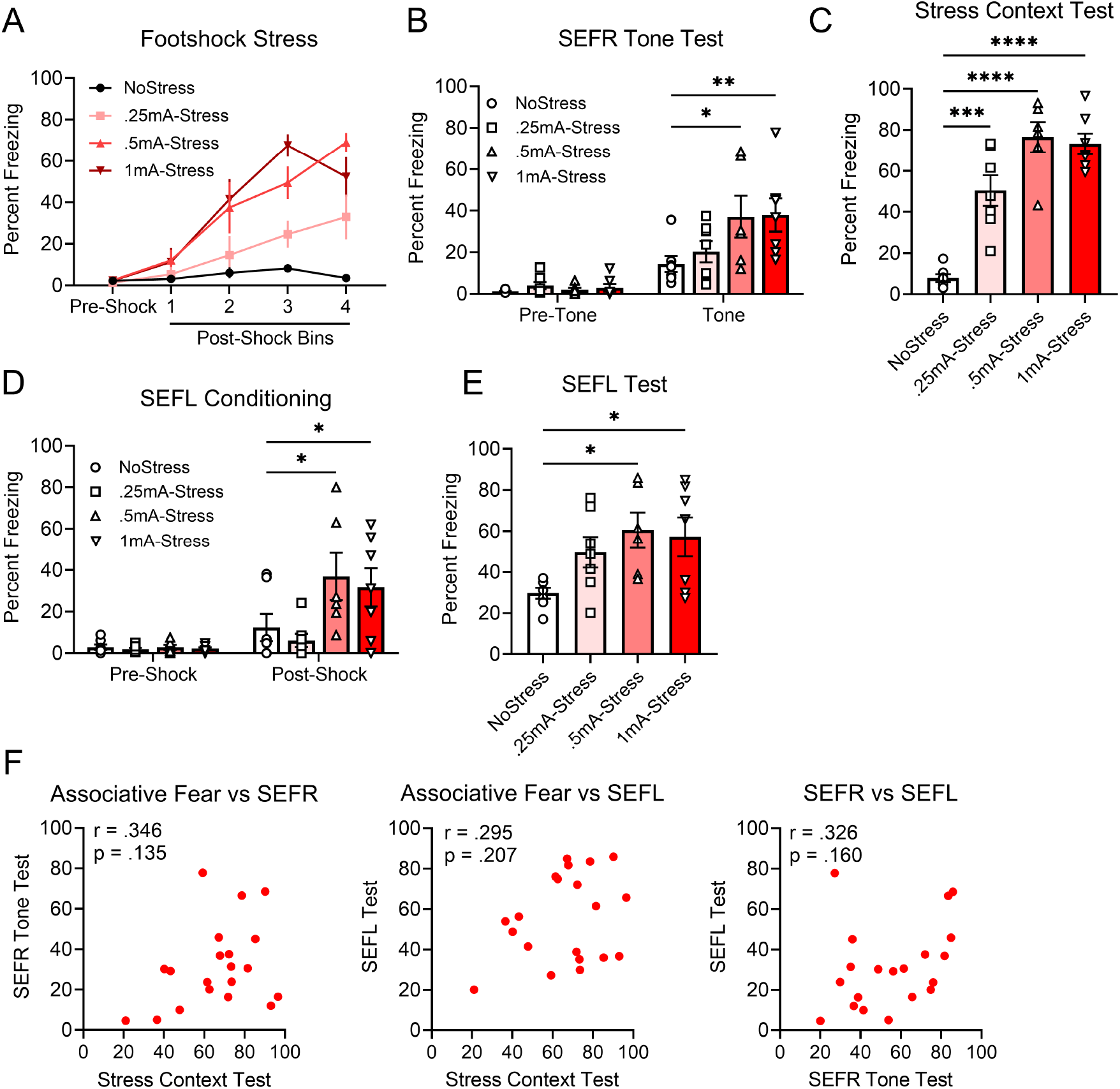
Varying the footshock stress intensity reveals dissociations among associative fear, SEFR, and SEFL. (A) Animals received either no footshocks (NoStress group) or footshocks at intensities of 0.25 mA, 0.5 mA, or 1 mA (Stress groups). In the stress session, all footshock stress-exposed groups exhibited elevated levels of freezing relative to the NoStress group. (B) In the SEFR tone test, the 0.5mA-Stress and 1mA-Stress groups showed significantly higher freezing than the NoStress group. (C) In the stress context test, the 0.25mA-Stress, 0.5mA-Stress, and 1mA-Stress groups displayed elevated freezing compared to the NoStress group. (D) During SEFL conditioning, no group differences in freezing were observed prior to the footshock. Following footshock, the 0.5mA-Stress and 1mA-Stress groups exhibited increased freezing relative to the NoStress group. (E) In the SEFL test, the 0.5mA-Stress and 1mA-Stress groups exhibited elevated freezing relative to the NoStress group. (F) Correlation of individual subjects’ percent freezing between the stress context test, SEFR tone test, and SEFL test, calculated among animals in the 0.25mA-Stress, 0.5mA-Stress, and 1mA-Stress groups. Data presented as mean +/-SEM. N=27: NoStress (7 males), 0.25mA-Stress (7 males), 0.5mA-Stress (6 males), 1mA-Stress (7 males). * *p* < 0.05, ** *p* < 0.01, *** *p* < 0.001.

### Stressed mice exhibit potentiated c-Fos expression in the paraventricular thalamus during SEFR

To identify brain regions that might mediate sensitization of unlearned fear, we examined the effects of prior footshock stress on c-Fos expression evoked during the SEFR tone test. On the first day of testing, mice underwent footshock stress (Stress groups) or an equal duration of context exposure (NoStress groups). Following 24 hrs, mice were placed in a novel context and underwent the SEFR tone test (ToneTest groups) or were left undisturbed in their homecage (HomeCage groups). After 90 min, brain tissue was collected and processed for immunohistochemistry against the c-Fos protein. This experimental design established the following groups: NoStress-HomeCage, Stress-HomeCage, NoStress-ToneTest, Stress-ToneTest.

Next, we quantified c-Fos expression across several limbic brain structures previously implicated in the regulation of fear and other defensive behaviors (Fig. 3A). A two-way ANOVA revealed a significant Group x Region interaction (*F*_18,202_ = 2.29, *p* <.0001), indicating that stress effects on neural activity varied across brain areas. Post hoc comparisons showed that relative to the NoStress-ToneTest group, the Stress-ToneTest group exhibited significantly elevated levels of c-Fos expression in the anterior paraventricular thalamus (aPVT) (*p* <.0005) and the posterior paraventricular thalamus (pPVT) (*p* <.0005). The c-fos levels in these groups did not differ in the basolateral amygdala (*p* =.9393), central amygdala (*p* =.9997), lateral parabrachial nucleus (*p* =.9917), dorsolateral periaqueductal gray (*p* =.5826), or lateral periaqueductal gray (*p* =.5356). c-fos levels in the NoStress-HomeCage and Stress-HomeCage groups did not differ significantly in any region (*p >*.05), indicating that stress did not alter baseline c-Fos expression.

**Figure 3.**
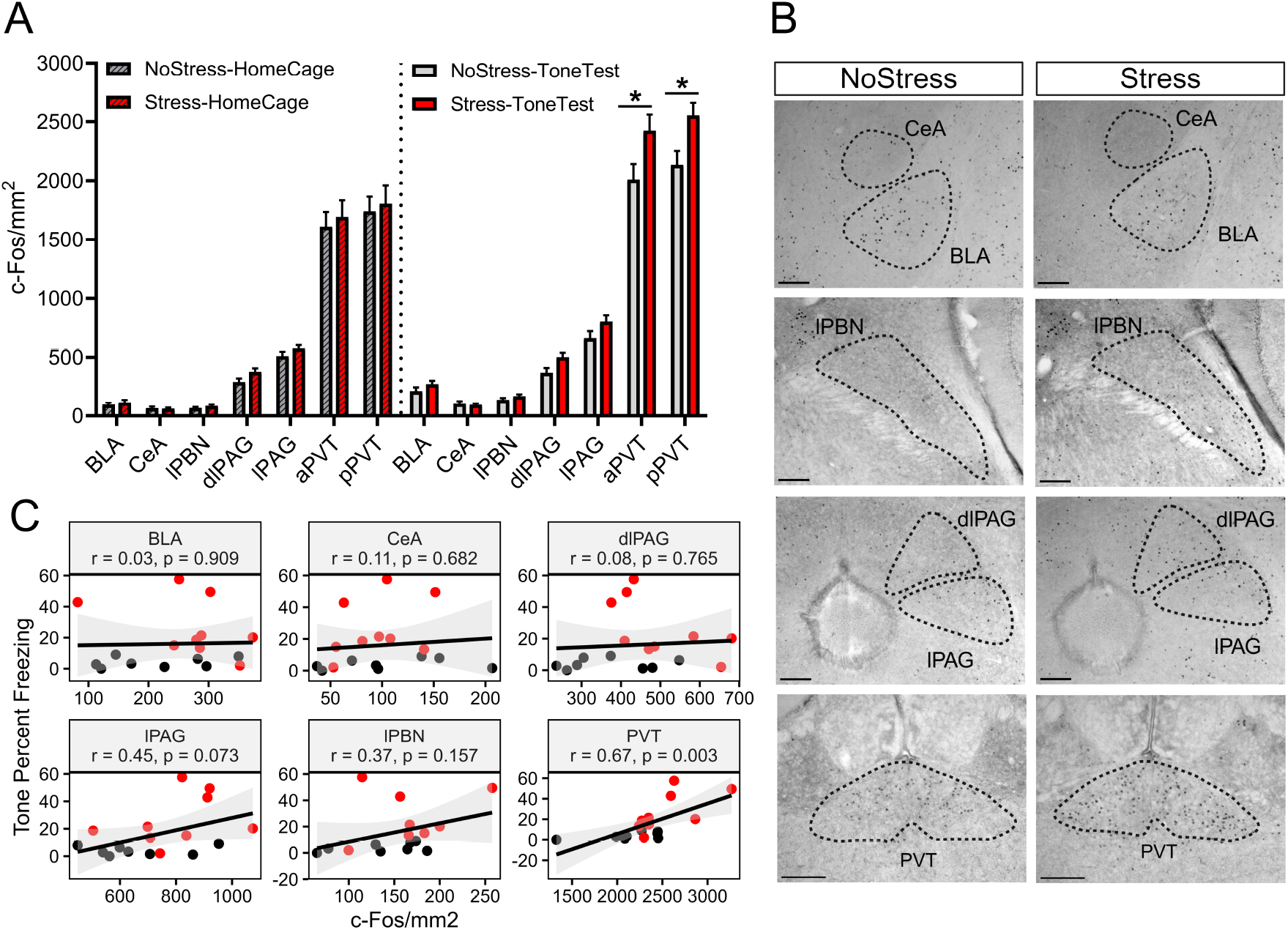
Footshock stress potentiates c-Fos expression in the paraventricular thalamus in response to the SEFR tone test. (A) Quantification of c-Fos expression across brain regions: BLA, basolateral amygdala; CeA, central amygdala; lPBN, lateral parabrachial nucleus; dlPAG, dorsolateral periaqueductal gray; lPAG, lateral periaqueductal gray; PVT, paraventricular thalamus. (B) Representative images of c-Fos expression in the NoStress-ToneTest and Stress-ToneTest groups. (C) Correlation between c-Fos expression and freezing levels during the tone period of the SEFR tone test. Black circles represent subjects in the NoStress-ToneTest group; red circles represent subjects in the Stress-ToneTest group. Data presented as mean +/-SEM. N=33: NoStress-HomeCage (8 males), Stress-HomeCage (8 males), NoStress-ToneTest (8 males), Stress-ToneTest (9 males) * *p* < 0.05, ** *p* < 0.01, *** *p* < 0.001.

These results demonstrate that footshock stress potentiates aPVT and pPVT activity in response to the SEFR tone test. Lastly, we examined the relation between c-Fos expression and freezing behavior during the SEFR tone test (Fig. 3C). c-Fos expression in the PVT (aPVT and pPVT average) was significantly correlated with freezing to the tone (*r* =.67, *p* =.003), whereas c-Fos expression in other regions did not correlate significantly. These findings highlight PVT activity as a potential regulator of sensitized unlearned fear responses.

### Stressed mice display elevated pPVT neural activity to a tone stimulus, but not to footshock

Our c-Fos analysis suggests that stressed mice display neuronal hyperactivity in both the aPVT and pPVT during the SEFR tone test. The PVT is a well-documented stress-responsive region activated by diverse aversive stimuli, such as foot shock, homeostatic deregulation, drug withdrawal, and social defeat (***Zhu et al., 2018***; ***Beas et al., 2018***; ***Dong et al., 2020***; ***Wang et al., 2021***). The PVT also regulates fear and defensive behavior via its extensive projections to limbic brain regions (***Penzo et al., 2015***; ***Padilla-Coreano et al., 2011***; ***Do-Monte et al., 2015a***; ***Ma et al., 2021***). For these reasons, the PVT is well-positioned to mediate stress effects on fear circuitry that may be important for fear sensitization. Although neurons throughout the PVT are activated by salient stimuli, evidence indicates that the pPVT exhibits greater responsiveness to aversive stimuli than the aPVT (***Choi et al., 2019***). Moreover, studies have identified long-term alterations in firing pattern and synaptic connectivity in the pPVT that modulate fear and defensive behaviors (***Do-Monte et al., 2015a***; ***Penzo et al., 2015***; ***Li et al., 2010***). For these reasons, we focused our investigation on the pPVT and sought to further examine its putative role in fear sensitization. We expressed the genetically encoded calcium indicator GCaMP8s in the pPVT and implanted an optical fiber above the region (Fig. 4A.) After the mice recovered, they were subjected to either footshock stress (Stress group) or context exposure with no footshocks (NoStress group). We then monitored bulk Ca^2^+ signals using fiber photometry during the SEFR tone test and SEFL conditioning session. During the recording sessions, individual freezing data could not be determined because the fiber optic patch cable interfered with mouse activity. The aim of this experiment was to confirm whether stress-induced alterations in pPVT activity are present during exposure to the novel tone stimulus used to evoke SEFR and/or the single mild footshock used to elicit SEFL.

**Figure 4.**
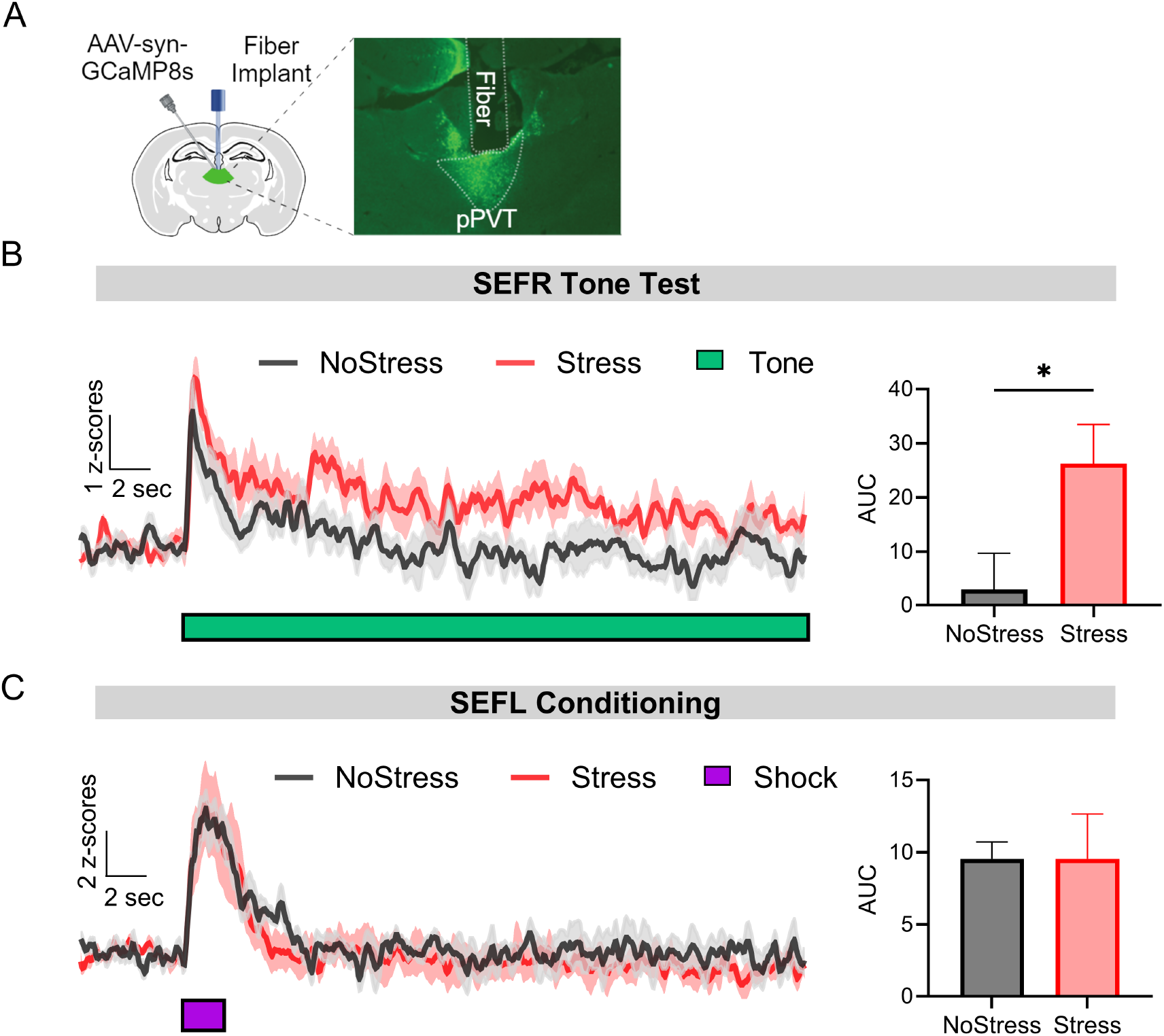
Mice previously exposed to footshock stress exhibit elevated pPVT activity in response to the novel tone used to elicit SEFR but not to the footshock used to induce SEFL. (A) Representative image GCamP8s expression and fiber track in the pPVT. (B) Left; Ca^2^+ signals of pPVT averaged across the three tone presentations in the SEFR tone test for the Stress and NoStress groups. Right; Quantification of AUC during the 30-s tone presentation compared between the NoStress and Stress groups. (C) Left; Ca^2^+ signals in pPVT during the SEFL conditioning session for the Stress and NoStress groups. Right; Quantification of AUC during the 2-s footshock presentation compared between the NoStress and Stress groups. Data presented as mean +/-SEM. For (B) N=16: NoStress group (6 males, 2 females), Stress group (6 males, 2 females). For (C) N=11: NoStress group (3 males, 2 females), Stress group (4 males, 2 females). * *p* < 0.05, ** *p* < 0.01, *** *p* < 0.001.

In the SEFR tone test, all mice exhibited Ca^2^+ signals in pPVT neurons that peaked shortly after tone onset and gradually declined over the 30-s stimulus (Fig 4B, left panel). Notably, the Stress group displayed higher levels of pPVT activity during the tone compared to the NoStress group (Fig. 4B, right panel; *t*_14_ = 2.358, *p* =.0335). We next examined pPVT activity responses to the footshock used in the SEFL conditioning session. Here, the presentation of the footshock evoked a rapid increase of Ca^2^+ signals that were comparable between the NoStress and Stress groups (Fig. 4C, left panel). AUC analysis during the 2-s presentation of the footshock revealed no significant group differences (Fig. 4C, right; (*t*_9_ =.00014, *p* =.9989). In summary, stress exposure enhanced pPVT Ca^2^+ responses in the SEFR tone test but did not alter pPVT responses during fear conditioning in a novel context. These results suggest that pPVT hyperactivity in stressed mice is a distinct feature of sensitization of unlearned fear but not sensitization of fear learning.

### Inhibition of pPVT during footshock stress disrupts the induction of SEFR but not SEFL

We next tested the hypothesis that pPVT neuronal activity during footshock stress is necessary for the induction of fear sensitization. An adeno-associated virus (AAV) encoding the inhibitory hM4Di receptor (AAV8-hSyn-hM4Di-mCherry) or the fluorescent mCherry reporter (AAV8-hSyn-mCherry) was infused in the pPVT of mice (Fig. 5A). Following recovery from surgery, we tested whether CNO-mediated inhibition of pPVT neurons during footshock stress would prevent subsequent stress-induced alterations in fear and anxiety-like behavior. All mice underwent the same behavioral testing procedure depicted in Fig. 5B.

**Figure 5.**
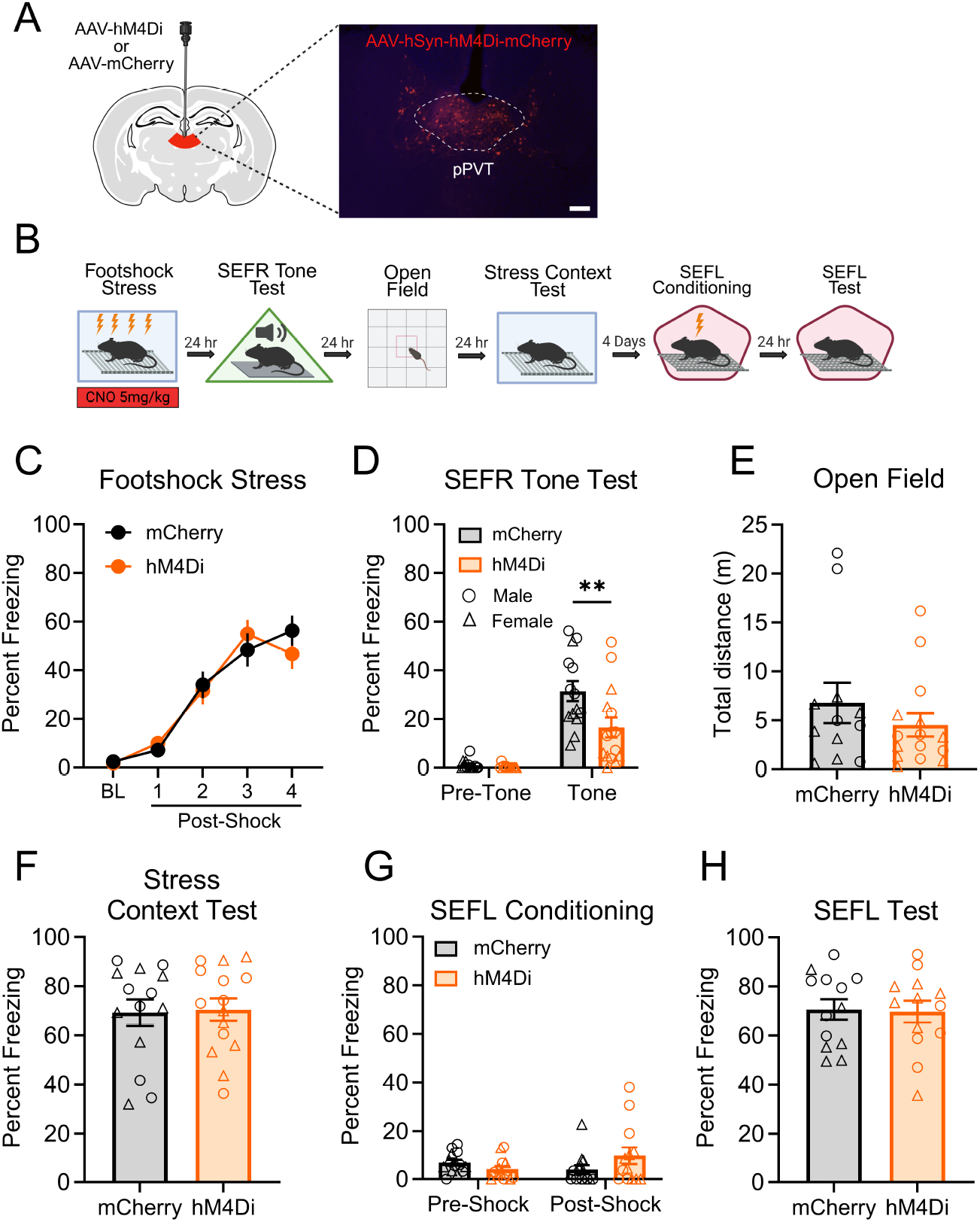
Inhibition of pPVT during footshock stress impaired the induction of SEFR but not SEFL. (A) Schematic of the AAV viral strategy and representative image of pPVT viral expression. Animals received AAVs encoding the inhibitory designer receptor Hm4Di or a control mCherry fluorescent reporter. Scale bar indicates 100 μm. (B) Experimental timeline. (C) Inhibition of pPVT had no effect on freezing behavior during footshock stress. (D) In the SEFR tone test, prior inhibition of pPVT during footshock stress decreased tone-elicited freezing. (E) The distance traveled in the open field test was comparable between the hM4Di and mCherry groups. (F) Associative fear of the stress context was not affected by pPVT inhibition during footshock stress. (G) In the SEFL conditioning session, mice displayed low levels of freezing prior to the presentation of footshock. (H) In the SEFL test session, all groups displayed similar levels of freezing to the context. Data presented as mean +/-SEM. N=29: Hm4Di group (8 males, 7 females), mCherry group (7 males, 7 females). * *p* < 0.05, ** *p* < 0.01, *** *p* < 0.001..

CNO was administered 30 min prior to footshock stress (Fig. 5C). During this session, freezing levels increased progressively as a function of shock number, but no significant differences between the groups were detected (*F*_1,27_ =.01788, *p* =.895). Subsequent behavioral assays were conducted without CNO administration. In the SEFR tone test (Fig. 5D), a RM-ANOVA revealed a significant Group x Tone Period interaction (*F*_1,27_ = 6.097, *p* =.0202). Freezing behavior was notably lower in the hM4Di group compared to the mCherry group. Post hoc tests confirmed that these differences in freezing were restricted to the tone period (*p* =.0013) and not the pre-tone baseline period (*p* =.9759).

Next, we conducted subsequent behavioral assays in the same cohort of mice to further investigate the role of pPVT in regulating additional stress-altered fear and anxiety-like behaviors. Our previous report showed that total distance traveled in the open field is reliably suppressed after footshock stress (***Hassien et al., 2020***). In the open field (Fig. 5E), the distance traveled was not significantly different between groups (*t*(25) =.9920, *p* =.3307). Time and distance in the center arena also did not differ between groups (data not shown). In the test for associative fear of the stress context (Fig. 5F), we also observed no significant group differences in freezing behavior (*t*(27) =.1735, *p* =.8636). During SEFL conditioning (Fig. 5G), we observed no significant main effect of Group (*F*_1,28_ =.4515, *p* =.5080). Lastly, in the SEFL test (Fig. 5H), expression of contextual fear did not differ between the mCherry and hM4Di groups (*t*(24) =.1506, *p* =.8815). Collectively, the results from this experiment demonstrate that inhibiting the pPVT during footshock stress attenuated SEFR but did not affect conditioning of the stress context, open field exploration, or SEFL.

### Inhibition of pPVT attenuates the expression of SEFR but not SEFL

The findings thus far suggest that pPVT activity is necessary for the induction of SEFR but not SEFL. Next, we asked whether pPVT activity is required for the expression of either of these forms of sensitization. As in the previous experiment, we used an AAV approach to express the inhibitory hM4Di receptor or the control mCherry fluorescent reporter in pPVT neurons (Fig. 6A). After recovery from surgery, all groups were exposed to footshock stress without CNO treatment. Subsequently, mice underwent behavioral testing under CNO treatment (CNO administered 30 min prior to each test) to assess the impact of pPVT inhibition on fear and anxiety-like behavior (Fig. 6B).

**Figure 6.**
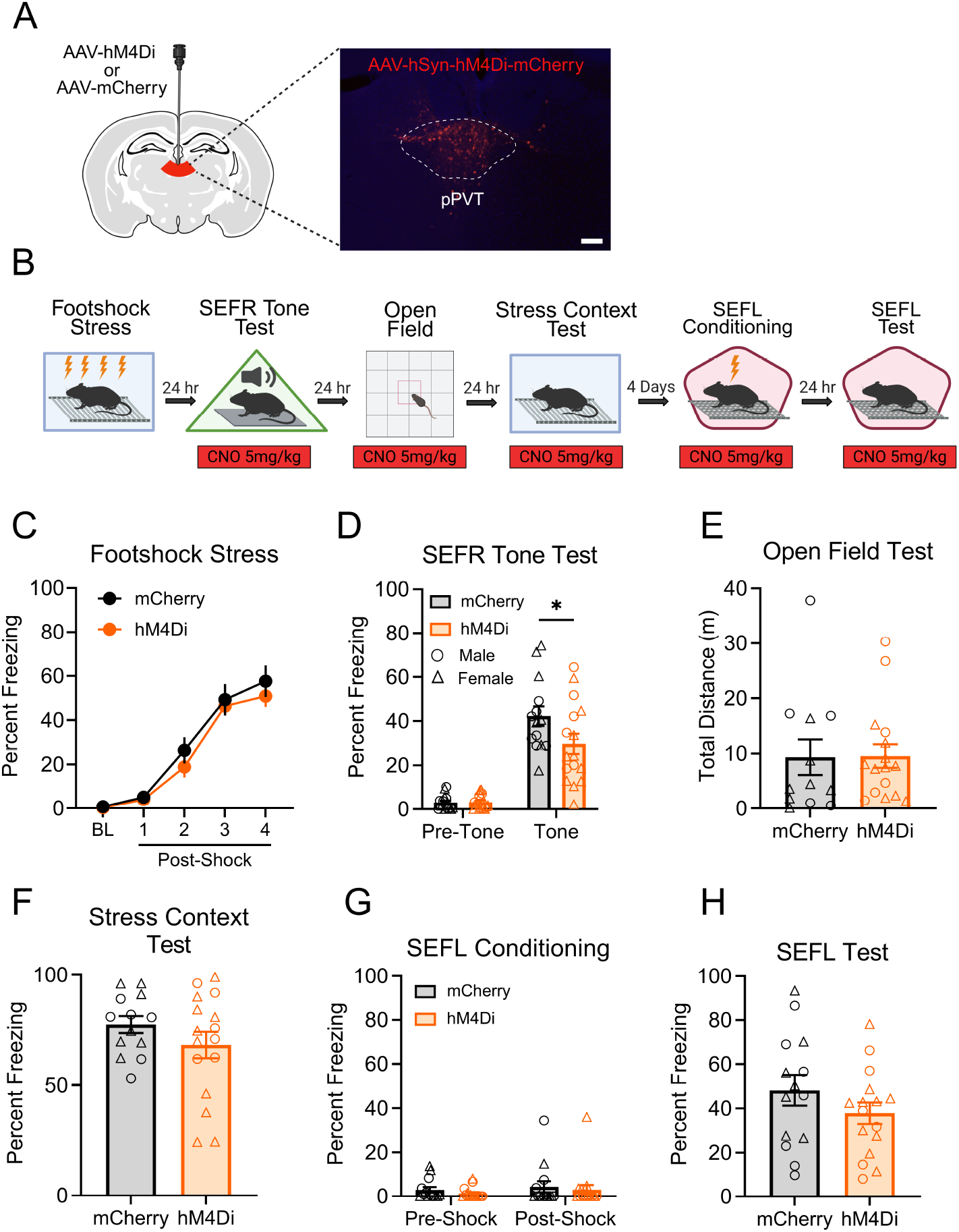
Inhibition of pPVT attenuates the expression of sensitization of unlearned fear but not sensitization of fear learning. (A) Schematic of the AAV viral strategy and representative image of viral expression in the pPVT. Mice received AAVs encoding the inhibitory Hm4Di receptor or the mCherry fluorescent reporter. Scale bar indicates 100 μm. (B) Experimental timeline. (C) In the footshock stress session, freezing behavior was comparable between the hM4Di and mCherry groups. (D) In the SEFR tone test, the hM4Di group displayed significantly decreased levels of freezing compared to the mCherry group, specifically during the tone period. (E) In the open field test, total distance traveled did not differ between the groups. (F) In the stress context test, both the hM4Di and mCherry groups showed similar levels of freezing. (G) In the SEFL conditioning session, mice displayed low levels of freezing prior to the footshock presentation. (H) In the SEFL test session, freezing behavior did not differ between the hM4Di and mCherry groups. Data presented as mean +/-SEM. N=30: Hm4Di group (6 males, 10 females), mCherry group (7 males, 7 females). * *p* < 0.05, ** *p* < 0.01, *** *p* < 0.001.

During the footshock stress session conducted in the absence of CNO (Fig. 6C), mice displayed progressively increasing levels of freezing in response to the footshocks. No significant differences were observed between groups that received the hM4Di or mCherry viral infusions (*F*_1,28_ =.8117, *p* =.3753). 24 hours later, mice underwent the SEFR tone test with CNO on board (Fig. 6D). A RM-ANOVA revealed a significant Group x Tone period interaction (*F*_1,28_ = 4.422, *p* =.0446). As confirmed by post hoc analysis, pre-tone baseline levels of freezing were similar between groups (*p* =.9995), but during the tone period, the hM4Di group displayed significantly less freezing than the mCherry group (*p* =.0178). These data suggest that CNO-mediated inhibition of pPVT neurons attenuated the behavioral expression of SEFR.

Next, we conducted the open field test with CNO on board (Fig. 6E). No significant differences in distance traveled or time spent in the center arena (data not shown) were observed between the hM4Di and mCherry groups (*t*_26_ =.0568, *p* =.9551), indicating no effect of pPVT inhibition on locomotor activity or anxiety-like behavior. We then examined the effect of pPVT inhibition on associative fear memory retrieval by measuring context-evoked fear of the footshock stress context with CNO on board (Fig. 6F). Here, freezing did not differ significantly between groups (*t*_27_ = 1.234, *p* =.2278). Lastly, we performed the two-day SEFL procedure with CNO administered during both the SEFL conditioning and SEFL test sessions. During SEFL conditioning (Fig. 6G), we observed no significant main effect of Group (*F*_1,28_ =.6567, *p* =.4246) or Group x Shock Period interaction (*F*_1,28_ =.0077, *p* =.9308). In the SEFL test session (Fig. 6H), freezing behavior did not differ significantly between groups (*t*_28_ = 1.245, *p* =.2234). Overall, CNO-mediated inhibition of pPVT did not affect anxiety-like behavior, expression of associative fear of the stress context, or SEFL. Notably, the effect of this manipulation on the SEFR tone test, but lack of an effect on SEFL, indicates that the pPVT selectively supports the expression of sensitization of unlearned fear but not the sensitization of fear learning.

### Artificial activation of pPVT in stress-naïve mice potentiates unlearned fear but not fear learning

Next, we asked whether chemogenetic activation of pPVT neurons in stress-naïve mice was sufficient to recapitulate the sensitized fear observed in footshock-stressed mice. An AAV encoding either the excitatory hM3Dq receptor (AAV8-hSyn-hM3Dq-mCherry) or the fluorescent mCherry reporter (AAV8-hSyn-mCherry) was infused into the pPVT (Fig. 7A). After recovery, mice were tested in the SEFR tone test, open field, and SEFL procedure. CNO was administered 30 min prior to each of the following behavioral sessions: the SEFR tone test, open field, and SEFL conditioning (Fig. 7B).

**Figure 7.**
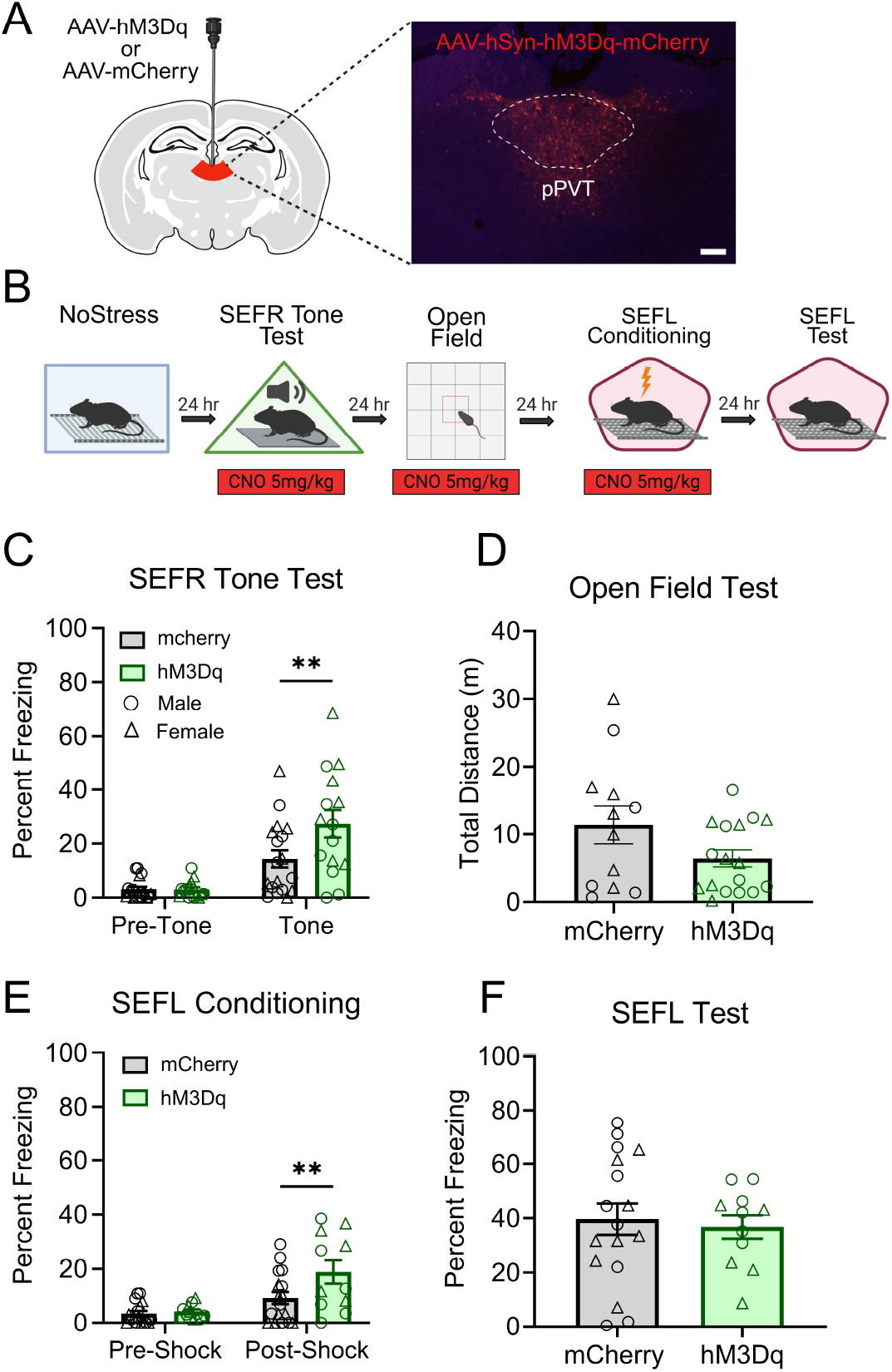
Activation of pPVT in stress-naïve mice potentiates freezing to the novel tone but does not enhance fear conditioning. (A) Schematic of the AAV viral strategy and representative image of viral expression in pPVT. Mice received AAVs encoding the excitatory Hm3Dq receptor or the mCherry fluorescent reporter. Scale bar indicates 100 μm. (B) Experimental timeline. (C) In the SEFR tone test, the Hm3Dq group displayed significantly higher levels of freezing compared to the mCherry group during the tone period. (D) In the open field, no significant group differences were observed in total distance traveled. (E) In the SEFL conditioning session, pre-Shock freezing levels did not differ between groups. However, the Hm3Dq group displayed increased post-shock freezing relative to the mCherry group. (F) In the SEFL test session, the hM3dq and mCherry groups showed similar levels of freezing. Data presented as mean +/-SEM. N=35: Hm3Dq (9 males, 8 females), mCherry (9 males, 9 females). * *p* < 0.05, ** *p* < 0.01, *** *p* < 0.001.

In the SEFR tone test (Fig. 7C), a RM-ANOVA revealed a significant Group x Tone Period interaction (*F*_1,31_ = 6.133, *p* =.0189). Post hoc analysis showed no group differences during the pretone baseline period (*p* =.9992). However, during the tone period, the hM3Dq group displayed significantly higher levels of freezing relative to the mCherry group (*p* =.0056). In the open field test (Fig. 7D), chemogenetic activation of the pPVT did not significantly affect distance traveled (*t*_27_ = 1.787, *p* =.0852). Finally, to examine whether activation of pPVT enhances fear learning to a novel context, we conduced the SEFL procedure with CNO administered prior to the SEFL conditioning session. During SEFL conditioning (Fig. 7E), a RM-ANOVA revealed a significant main effect of Shock Period (*F*_1,26_ = 31.12, *p* <.0001) and a significant Group x Shock Period interaction (*F*_1,26_ = 5.939, *p* =.0220). Pre-shock freezing levels did not differ between groups, indicating that CNO administration did not alter baseline fear. However, following footshock, the hM3Dq group displayed significantly elevated freezing compared to the mCherry group (*p* =.0099). Despite this, freezing levels in the SEFL test session did not differ between groups (Fig. 7F; *t*_26_ = 0.3591, *p* =.7224). In summary, chemogenetic activation of the pPVT in stress-naïve mice was sufficient to phenocopy sensitization of unlearned fear to a novel tone (SEFR), but did not enhance fear learning (SEFL).

### Inhibition of pPVT fails to attenuate spontaneous tone-elicited fear in stress-naïve mice

Our results indicate that pPVT neuronal activity is required for the sensitizing effects of stress in the SEFR tone test. However, it is unclear whether PVT activity modulates fear expression generally or only after exposure to footshock stress. To test whether chemogenetic inhibition of pPVT could diminish freezing behavior in the SEFR tone test with stress-naïve mice, we conducted a tone test in which the tone amplitude was increased to a level (115 dB) at which it evokes freezing even in stress-naive mice. Mice received viral infusion of AAV encoding the inhibitory hM4Di receptor or the fluorescent mCherry reporter (Fig. 8A). Following recovery, all mice were tested in the modified SEFR tone test with CNO on board (Fig. 8B). A RM-ANOVA on freezing behavior revealed a main effect of Tone Period (*F*_1,17_ = 37.50, *p* <.0001), demonstrating that the tone was of sufficient intensity to elicit elevated levels of freezing relative to the pre-tone period. However, we did not detect a significant main effect of Group (*F*_1,17_ =.07030, *p* =.7941) or a significant Group x Tone Period interaction (*F*_1,17_ =.06350, *p* =.8041). This finding suggests that pPVT neuronal activity does not modulate expression of fear in mice that have not been exposed to stress.

**Figure 8.**
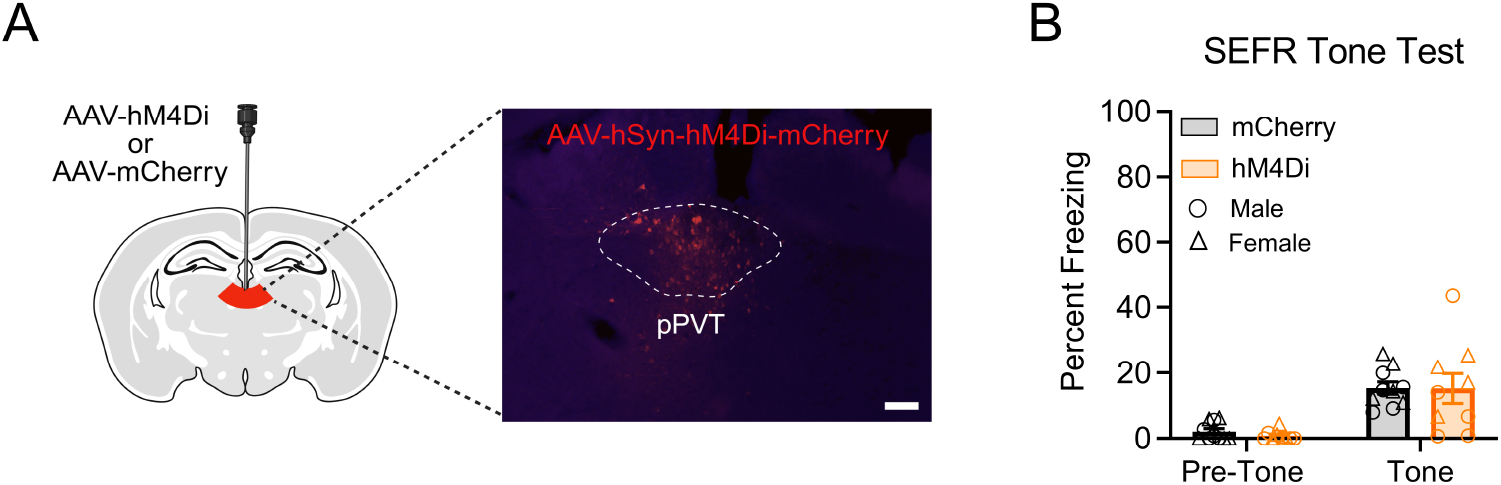
Inhibition of pPVT in stress-naïve mice does not attenuate freezing responses to a 115-dB tone. (A) Schematic of the AAV viral strategy and representative image of viral expression in pPVT. Animals received an AAV encoding the inhibitory hM4Di receptor or a fluorescent mCherry reporter. Scale bar indicates 100 μm. (B) In the SEFR tone test, no significant group differences in freezing behavior was observed. Data presented as mean +/-SEM. N=19: Hm4Di (5 males, 4 females), mCherry (5 males, 5 females). * *p* < 0.05, ** *p* < 0.01, *** *p* < 0.001.

## Discussion

In this study, we identified a novel role for the pPVT in stress-induced fear sensitization. We found that brief exposure to footshock stress causes long-lasting hyperexcitability of the pPVT. This hyperactivity was selective: it was detected in response to a novel tone used to demonstrate SEFR but not to the footshock used to induce SEFL. Using chemogenetic inhibition, we demonstrated that pPVT activity is necessary for both the induction and expression of SEFR, but not SEFL. Conversely, artificial activation of the pPVT mimicked the effects of footshock in the SEFR tone test but failed to enhance fear conditioning in a novel context. Together, these findings suggest that pPVT hyperactivity mediates SEFR but not SEFL.

Stress is known to sensitize a range of fear and defensive behaviors, but few studies have directly tested whether different forms of fear sensitization arise from a shared mechanism or are mediated by behavior-specific circuits. Recent work suggests the latter. Manipulations of the dorsal hippocampus or ventromedial prefrontal cortex affect cued and contextual SEFL differently (***Hersman et al., 2019***; ***Pennington et al., 2017***). Moreover, ***Pennington et al. (2024***) showed that foot-shock stress-induced alterations in SEFL and avoidance behavior are selectively mediated by the basolateral amygdala and ventral hippocampus, respectively. Our results support this emerging view, revealing that SEFR and SEFL are dissociable forms of fear sensitization governed by distinct neural substrates.

Although numerous studies have used the SEFL procedure to investigate the nonassociative-related symptoms of PTSD (***Nishimura et al., 2022***), less is known about the behavioral phenotype observed in SEFR. We posit that the tone used in the SEFR test represents an innocuous stimulus of ambiguous threat potential. Although we observed little to no tone-elicited freezing in non-stressed mice, a study by ***Kamprath and Wotjak*** (***2004***) found that a novel tone could trigger freezing responses in non-stressed mice at decibel levels exceeding those used in our procedure (*>*90db). Thus, in our mouse model, the tone-elicited freezing in stressed mice likely reflects a left-ward shift (reduced threshold) in the expression of unconditioned fear. Additionally, SEFR is distinct from other behavioral assays of hyperarousal because freezing behavior is the primary index used for evaluation of unconditioned fear responses. In rodents, hyperarousal is commonly assessed by measuring startle reactions to loud, abrupt auditory stimuli (***Davis, 1989***). Additionally, procedures that measure a subject’s propensity to interact with innately fearful or threatening stimuli are also used as measures of hyperarousal and are thought to capture the hypervigilant or anxiety-like aspects of behavior. These tests often involve measuring behaviors such as avoidance or decreased exploration in procedures such as the open field, elevated plus maze, or predator odor interaction. However, these tests may not actually reflect nonassociative fear sensitization. Indeed, previous work from our lab has shown that decreased exploratory behavior in the open field following foot-shock stress is ameliorated by extinction of the stress context (***Hassien et al., 2020***), suggesting that aspects of associative fear memory mediate this phenotype. Taken together, we suggest that SEFR provides unique insight into the symptoms of PTSD that may relate to hyperarousal.

Although silencing the pPVT in stressed mice produced robust effects on SEFR, the same manipulation did not affect associative fear of the stress context or SEFL. These findings were somewhat unexpected, given prior studies showing that the PVT modulates both the acquisition and expression of Pavlovian fear conditioning (***Penzo et al., 2015***). However, the literature on this topic is mixed. While some studies demonstrate a role for the PVT in associative fear learning (***Penzo et al., 2015***; ***Do-Monte et al., 2015a***), others have reported selective or no effects following PVT lesions or inactivation (***Li et al., 2014***; ***Choi et al., 2019***). These inconsistencies could be explained by the functional heterogeneity of the PVT (***McGinty and Otis, 2020***). Distinct PVT cell populations can exert opposing effects on behavior (***Ma et al., 2021***), highlighting a potential limitation of global silencing, which may obscure effects of behaviorally relevant subpopulations. Moreover, several reports suggest a time-dependent role for the PVT in supporting associative fear memories, in which the PVT is engaged during the retrieval of late, but not early, time points (***Quiñones-Laracuente et al., 2021***; ***Do-Monte et al., 2015b***). In the present study, manipulations to the PVT were conducted at fixed times and it remains unclear whether the PVT supports SEFR in a similar time-dependent manner. This possibility warrants further investigation.

Our observation that the pPVT contributes to SEFR is consistent with recent evidence that stress inclines animals towards heightened levels of fear when faced with uncertain or ambiguous threats and that the PVT mediates this stress-induced shift in threat evaluation (***Wang et al., 2023, 2024***). In our study, the SEFR procedure involves presenting a novel tone in a novel chamber to mice with no prior experience with the stimulus or context. This makes the tone stimulus, although loud, inherently ambiguous in terms of its predictive value for danger. We suspect that the ambiguous nature of the tone engages the pPVT, and stress-induced hyperactivation of this region biases mice towards increased fear responses to such stimuli. This interpretation may also explain the lack of pPVT involvement in SEFL, where the explicit pairing of a context with footshock makes the threat value of the context unambiguous.

Our findings identify pPVT hyperactivity as a mechanism by which stress sensitizes unlearned fear. This interpretation aligns with prior reports showing that acute stress primes PVT responses to subsequent threats (***Pliota et al., 2018***; ***Beas et al., 2018***; ***Jász et al., 2025***). One potential driver of this hyperactivity is stress-induced plasticity within the pPVT. Notably, the PVT is tightly regulated by stress hormones, neurotransmitters, and neuropeptides (***Li and Kirouac, 2012***; ***Bentivoglio et al., 1991***; ***Curtis et al., 2021***). These modulatory signals may potentiate PVT activity through alterations in intrinsic neuronal properties. For example, stress-related neuromodulators such as dynorphin (***Chen et al., 2015***) and orexins (***Kolaj et al., 2007***) are known to influence cellular excitability in the PVT. Alternatively, stress may alter presynaptic inputs to the PVT to facilitate increased activity. For instance, dopamine released from locus coeruleus terminals during footshock stress suppresses inhibitory pre-synaptic input onto PVT neurons, thereby increasing their excitability (***Beas et al., 2018***).

Our study shows that pPVT is necessary for the induction of SEFR during stress, suggesting that it receives and processes information about the stressor. Identifying the specific projection pathways that transmit stress information to the pPVT is crucial for understanding how stress initiates the sensitization process. Anatomical studies point to brainstem inputs like the lateral parabrachial nucleus (lPBN) and dorsolateral periaqueductal gray (dlPAG) as potential sources of nociceptive information projecting to pPVT (***Li and Kirouac, 2012***). Both lPBN and dlPAG are involved in transmitting threat signals and modulating defensive behavior (***Li and Kirouac, 2012***; ***Sato et al., 2015***; ***Kim et al., 2013***). Future studies should leverage techniques like viral-mediated retrograde tracing to identify specific neuronal populations projecting to pPVT that are activated by aversive stimuli and contribute to SEFR induction.

At present, it remains unclear whether specific populations of pPVT neurons are responsible for both the induction and expression of fear sensitization. Neurons in the PVT exhibit significant heterogeneity based on distinct gene expression, activity patterns, and circuit connectivity (***Kirouac, 2015***). Moreover, unique cell types have been shown to exert opposing influences on behavior, with distinct “stress-responsive” populations identified (***Gao et al., 2020***; ***Beas et al., 2018***). The possibility that stressful experiences are “encoded” by a subset of pPVT neurons and subsequently reactivated by ambiguous threats to sensitize fear responses is intriguing. A recent study using activity-tagging demonstrated that PVT ensembles active during early life stress regulate adult behavior (***Kooiker et al., 2024***), supporting the idea of stress-specific memory traces. Future characterization of pPVT heterogeneity using modern techniques like single-nucleus RNA-sequencing is a critical next step to clarify whether distinct neuronal subpopulations underlie the induction and expression of SEFR.

The PVT has been proposed to integrate prior aversive experiences and regulate affective behaviors through its widespread limbic connections. Supporting this role, previous studies have shown that PVT activity contributes to stress-induced alterations in anxiety-like behavior, passive coping, and avoidance (***Zhu et al., 2022***; ***Pliota et al., 2018***; ***Ma et al., 2021***; ***Dong et al., 2020***). However, these studies often do not clearly distinguish whether such changes arise from associative or nonassociative processes. For instance, some stress-induced behavioral alterations can be reversed by extinction or through pharmacological suppression of the stress memory, suggesting that associative mechanisms may be involved (***Hassien et al., 2020***; ***Blundell et al., 2005***). To clarify the specific contribution of the pPVT to nonassociative fear sensitization, we demonstrated that it is essential for SEFR, which is a nonassociative behavioral phenotype. These findings expand the known behavioral functions of the PVT and position the pPVT as a key node in the circuitry linking stress to heightened unlearned fear responses.

In conclusion, our study characterized a set of behavioral and neural mechanisms by which exposure to stress leads to the induction and expression of sensitization of unlearned fear (SEFR). We established that SEFR and SEFL are distinct behavioral manifestations of fear sensitization supported by discrete neural mechanisms. Specifically, we found that exposure to footshock stress leads to long-lasting alterations in pPVT neural activity patterns that selectively underlie SEFR. These results have important implications for PTSD, particularly for symptoms such as hyperarousal and reactivity, which our SEFR tone test is intended to model. This work emphasizes the need for continued research into the nonassociative consequences of stress to achieve a comprehensive understanding of the full range of PTSD symptoms. Our findings highlight the pPVT as a potential therapeutic target for stress-induced sensitization of unlearned fear, a class of symptoms that may be resistant to therapies focused solely on associative processes.

## Methods and Materials

### Subjects

Adult male and female mice (12-18 weeks) from 129S6/SvEvTac background (Taconic, Germantown, NY) were bred in house, weaned at postnatal day 21 and group housed (3-5 per cage). Mice were maintained on a 12h light/dark cycle with food and water available ad libitum. Experiments were conducted during the light phase. Mice were randomly assigned to groups prior to each experiment. Since no significant sex-by-group interactions were observed, data from males and females were combined for analysis. All animal procedures were conducted according to the National Institutes of Health guidelines on the Care and Use of Laboratory Animals and were approved by the University of Texas at Austin Institutional Animal Care and Use Committee.

### Apparatus

Conditioning chambers (30.5 × 24 × 21) consisted of three aluminum walls and ceiling, a clear Plexiglas door, and stainless-steel grid flooring (Med Associates VFC-005A). Chambers were individually housed in sound-attenuating cabinets and illuminated with overhead white light throughout the entire session. The conditioning chambers were equipped with near-infrared cameras that recorded at 30 frames/s and a speaker for tone delivery. Videos were automatically scored (Video Freeze software; Med Associates) using a linear pixel change algorithm that defined freezing as the absence of all movement, except for those related to breathing. In order to design distinct contextual environments across behavioral procedures (Footshock stress, SEFR tone test, and SEFL procedure), modifications were made to the chamber’s scent, floor, walls, ceiling, and transportation.

Open field apparatus consisted of a 40 x 40 cm^2^ arena with opaque plastic walls 35 cm high. The arena was illuminated with white incandescent bulbs producing 85 lux at the center of the arena. Behavior was recorded using digital video cameras mounted above the arenas and analyzed using AnyMaze software (Stoelting Inc.).

### Behavioral procedures

Footshock stress: Mice received four footshocks (2 sec duration, 1.0 mA intensity), delivered through the grid floor of the chamber at 160, 240, 320, and 400 sec after placement. Forty seconds following the final footshock, mice were returned to their homecage. NoStress control mice were exposed to the chamber for the same duration (7 min total) but did not receive any shocks. To promote contextual discrimination between the stress context and future testing environments, mice underwent extinction sessions consisting of 5-minute re-exposures to the footshock context once daily for five consecutive days following the stress session. The first extinction session also served as the test for stress context recall.

SEFR tone test: Following a 3 min acclimation period, mice were presented with three 30 sec tones (90 dB, 9 kHz, 50 ms rise/fall time), each separated by a 30 sec interstimulus interval. For analysis, freezing behavior was segmented into two epochs: the pre-tone period (defined as the duration before the onset of the first tone) and the tone period (defined as the duration following the onset of the first tone). The tone period includes freezing during all three tone presentations and the corresponding interstimulus intervals, as freezing behavior was consistent across these events. Freezing behavior for each epoch was averaged because we did not detect significant differences across time. An earlier study from our group reported that female, but not male, mice exhibited elevated freezing to the tone regardless of prior stress exposure (***Hassien et al., 2020***). Since that publication, the protocol has been refined to eliminate this effect. Specifically, we increased the discriminability between the footshock stress and SEFR tone test contexts by modifying visual, olfactory, and transportation cues. As a result, we no longer observe enhanced freezing in stress-naïve female mice under the updated protocol.

Open Field: Testing was conducted in a separate behavioral room from the other behavioral experiments. Mice were allowed to explore the arena for 30 min.

Stress-enhanced fear learning: On Day 1 of the SEFL procedure (SEFL conditioning session), mice received a single footshock (2 sec, 0.75 mA) delivered through the grid flooring 3 min after the start of the session. Mice were removed from the chambers 30 sec after footshock delivery and returned to homecage. On day 2 (SEFL test session), mice were placed into the same context from the previous day for 5 min to assess context-elicited freezing.

### Surgery

Mice were anesthetized using isoflurane (3% in induction chamber, maintained at 1.5% for remainder of surgery) and were placed in a stereotaxic apparatus. Prior to skin incision to reveal the skull, mice received a subcutaneous injection of buprenorphine (10 mg/kg) and carprofen (0.1mg/kg). Body temperature was maintained at 37 °C with a heating pad and ophthalmic ointment was applied to prevent the eyes from drying. The skull was exposed and a 1 mm diameter craniotomy was made at the site of injection. A pulled glass containing virus was slowly lowered to the target site. Viral coordinates were selected according to the atlas of Paxinos and Franklin. After viral infusions, the injector remained in place for 10 min to allow for viral diffusion.

For chemogenetic manipulation experiments, mice received unilateral viral infusion of AAV8-hSyn-hM3Dq-mCherry (Addgene: 50474), AAV8-hSyn-hM4Di-mCherry (Addgene: 50475), or AAV8-hSyn-mCherry (Addgene: 114472) in the pPVT (AP −1.6, ML 0.0, DV −3.3). Virus was delivered at a rate of 23.5nL/min for a total of amount of 138 nL. Following 3-4 weeks of recovery, clozapine-N-oxide (5mg/kg) was administered via I.P. injection 30 min prior to the start of behavior testing.

For fiber photometry experiments, mice received unilateral viral infusion of AAV1-syn-jGCaMP8s-WPRE (Addgene: 162374) in pPVT (AP −1.6, ML 0.0, DV −3.3). Following viral infusion, a second craniotomy was made and acidic gel etchent was applied to the surface of the skull followed by a layer of OptiBond epoxy (Kerr Corporation). Optical fiber cannulas with 400 um core diameter (Doric Lenses) were unilaterally implanted at a 7 degree angle targeting the pPVT (AP −1.6, ML 0.0, DV −3.3). Dental cement (Bosworth) was applied to secure the cannula to the skull. Mice were given 2 weeks to recover prior to photometry recordings.

### Tissue preparation and histology

Mice were deeply anesthetized with ketamine/xylazine (150/15mg/kg) and transcardially perfused with 0.01M phosphate buffered saline (1x PBS), followed by 4% paraformaldehyde (PFA) in 1x PBS. Brains were extracted and post-fixed at 4C in 4% PFA overnight then transferred to 30% sucrose solution in 1x PBS at 4C for 24 hr prior to cryopreservation. Coronal tissue sections (35um) were prepared on a cryostat.

For c-Fos immunohistochemistry, sectioned tissue was incubated in 1x PBS/methanol solution (1:1) with 1% hydrogen peroxide for 15 min. Sections were then washed in 1x PBS with 0.5% Triton-X (1x PBST) three times and blocked at room temperature in 1x PBST with 5% normal donkey serum (NDS) for 1.5 hr. Sectioned tissue was then incubated in for 24 hr at 4C degrees in a primary antibody solution (1:2000 rabbit anti-c-Fos polyclonal; Synaptic Systems 226-003) diluted in 1x PBST with 5% NDS. Sections were then washed in 1x PBST and then treated with secondary antibody solution (biotinylated goat anti-rabbit 1:500; Abcam ab64256) for 2 hrs at room temperature. Tissue sections were washed in 1x PBS then incubated in Vectastain Elite ABC reagent for 30 min followed by 1x PBS wash. Next, tissue sections were incubated in DAB peroxidase substrate solution (Vector laboratories) for 8 min then immediately washed in 1x PBS and mounted onto slides. Lastly, tissue sections were dehydrated through a graded series of alcohol concentrations, cleared with Citrasolv (Fisher Scientific), and coverslipped with Fluoromount-G (SouthernBiotech).

For fiber photometry and chemogenetic manipulation experiments, brain tissue was collected at the conclusion of behavioral testing. Tissue sections were stained with 1:1000 DAPI solution and mounted onto slides before being coverslipped with Fluoromount-G (SouthernBiotech). Fluorescent images were obtained using a Zeiss Axio Zoom.V16. Viral expression and optical fiber placement were confirmed according to the mouse brain atlas of Franklin and Paxinos.

### Cell counting and quantification

Images of c-Fos positive cells were obtained using a Zeis Axio Lab.A1 microscope and Zen imaging software. Images were collected at the same exposure settings and dimensions. Regions of interest were chosen a priori and a sample size of approximately 3-5 images per region were collected for each subject. Images were quantified in ImageJ using an automated counting algorithm based on experimenter-defined cut off metrics including the size, circularity, and signal intensity relative to background. Automated cell quantification was validated by comparing results to manual counts in a randomly selected subset of images. The c-Fos quantification was calculated by averaging the number of c-Fos positive cells per mm2, per animals, then per group.

### Fiber photometry

in vivo recordings of fluorescent signal were collected using a commercial fiber photometry system (Tucker-Davis Technologies; RZ10x) and data acquisition was performed on TDT synapse software. Excitation of GCaMP8s and isosbestic signals was achieved using blue 465 nm (Lx465; TDT) and ultraviolet 405 nm (Lx405) LEDs sinusoidally modulated at 331 Hz and 211 Hz, respectively. The mean fiber power was 40-60*μ*W. Excitation light was passed through a dichroic mirror (Doric Minicube) coupled to a fiber optic patch cord (400 um fiber core diameter, 0.66 NA; Doric Lenses) that was connected to the cranial implanted fiber (200 um fiber core diameter, 0.66 NA; Doric Lenses). Emission signals were filtered at 6 Hz and acquired at a sampling frequency of 1017 Hz. Fiber photometry data were analyzed using custom-written MATLAB scripts adapted from ***Barker et al. (2017***). To account for time-dependent slow drift artifacts caused by photobleaching, a least-squares linear fit was applied to the isosbestic signal to obtain a fitted control signal. *dF* /*F* was computed by subtracting the fitted control signal (*F*_0_) from the 465 nm raw signal (*F*), and dividing by the fitted control signal (*F* − *F*_0_/*F*_0_). For epoch-based analysis, Z-scores were calculated as (*dF* /*F* − *dF* /*F*_*mean*_)/*dF* /*F*_*SD*_) where the mean and standard deviation correspond to the 5 seconds prior to each event of interest.

### Statistical analysis

For freezing behavior analysis, we calculated the percentage of time mice spent freezing during predefined behavioral event windows. For the footshock stress session, freezing was quantified during the pre-shock baseline period and each of the four post-shock periods. For the SEFR tone test, if no significant differences were observed between the three tone presentations and the subsequent interstimulus intervals, freezing was calculated across two time windows: the pre-tone period (time prior to the first tone) and the tone period (time following the onset of the first tone, including all tones and intervals). For the SEFL conditioning session, freezing was measured during the pre-shock baseline and the 30 sec post-shock period prior to removal from the chamber. Lastly, for the stress context test and SEFL test, the percentage of time spent freezing was calculated for the entire 5 min test sessions. For both the SEFR and SEFL tests, animals that exhibited *>*15 percent freezing during the SEFL conditioning pre-shock period or the SEFR pre-tone period were excluded from analysis. This pre-determined criteria was established prior to the experiments to minimize the confound of fear generalization observed in a small set of animals.

Freezing data were analyzed using ANOVA or repeated-measures ANOVA when appropriate. Significant main effects and interactions were probed using Sidak’s post hoc tests for between-subject pairwise comparisons and Bonferroni-adjusted paired t-tests for within-subjects comparisons, except for Fig. 2 in which Dunnett’s test was used to compare multiple stress groups against a single NoStress group. For c-fos analysis, data were analyzed using a two-way ANOVA (Brain Region X Group) with Tukey’s-corrected post hoc tests. Statistical analyses were performed in Prism 6 (GraphPad Software).

## Acknowledgments

Supported by NIH grants R01MH117426 and R21MH128610 to MRD, T32MH106454 to KJN.

## References

Barker, D. J., Miranda-Barrientos, J., Zhang, S., Root, D. H., Wang, H.-L., Liu, B., Calipari, E. S., and Morales, M. (2017). Lateral preoptic control of the lateral habenula through convergent glutamate and gaba transmission. Cell reports, 21(7):1757–1769.

Beas, B. S., Wright, B. J., Skirzewski, M., Leng, Y., Hyun, J. H., Koita, O., Ringelberg, N., Kwon, H.-B., Buonanno, A., and Penzo, M. A. (2018). The locus coeruleus drives disinhibition in the midline thalamus via a dopaminergic mechanism. Nature neuroscience, 21(7):963–973.

Bentivoglio, M., Balercia, G., and Kruger, L. (1991). The specificity of the nonspecific thalamus: the midline nuclei. Progress in brain research, 87:53–80.

Bienvenu, T. C., Dejean, C., Jercog, D., Aouizerate, B., Lemoine, M., and Herry, C. (2021). The advent of fear conditioning as an animal model of post-traumatic stress disorder: Learning from the past to shape the future of ptsd research. Neuron, 109(15):2380–2397.

Blundell, J., Adamec, R., and Burton, P. (2005). Role of NMDA receptors in the syndrome of behavioral changes produced by predator stress. Physiology & Behavior, 86(1–2):233–243.

Charney, D. S., Deutch, A. Y., Krystal, J. H., Southwick, S. M., and Davis, M. (1993). Psychobiologic Mechanisms of Posttraumatic Stress Disorder. Archives of General Psychiatry, 50(4):294–305.

Chen, Z., Tang, Y., Tao, H., Li, C., Zhang, X., and Liu, Y. (2015). Dynorphin activation of kappa opioid receptor reduces neuronal excitability in the paraventricular nucleus of mouse thalamus. Neuropharmacology, 97:259–269.

Choi, E. A., Jean-Richard-dit Bressel, P., Clifford, C. W. G., and McNally, G. P. (2019). Paraventricular Thalamus Controls Behavior during Motivational Conflict. Journal of Neuroscience, 39(25):4945–4958.

Curtis, G. R., Oakes, K., and Barson, J. R. (2021). Expression and Distribution of Neuropeptide-Expressing Cells Throughout the Rodent Paraventricular Nucleus of the Thalamus. Frontiers in Behavioral Neuroscience, 14:634163.

Davis, M. (1989). Sensitization of the acoustic startle reflex by footshock. Behavioral Neuroscience, 103(3):495.

Daws, S. E., Jamieson, S., de Nijs, L., Jones, M., Snijders, C., Klengel, T., Joseph, N. F., Krauskopf, J., Kleinjans, J., Vinkers, C. H., et al. (2020). Microrna regulation of persistent stress-enhanced memory: This article has been corrected since advance online publication and a correction is also printed in this issue. Molecular psychiatry, 25(5):965–976.

Do-Monte, F. H., Quiñones-Laracuente, K., and Quirk, G. J. (2015a). A temporal shift in the circuits mediating retrieval of fear memory. Nature, 519(7544):460–463.

Do-Monte, F. H., Quiñones-Laracuente, K., and Quirk, G. J. (2015b). A temporal shift in the circuits mediating retrieval of fear memory. Nature, 519(7544):460–463.

Dong, X., Li, S., and Kirouac, G. J. (2020). A projection from the paraventricular nucleus of the thalamus to the shell of the nucleus accumbens contributes to footshock stress-induced social avoidance. Neurobiology of Stress, 13:100266.

Gao, C., Leng, Y., Ma, J., Rooke, V., Rodriguez-Gonzalez, S., Ramakrishnan, C., Deisseroth, K., and Penzo, M. A. (2020). Two genetically, anatomically and functionally distinct cell types segregate across anteroposterior axis of paraventricular thalamus. Nature Neuroscience, pages 1–12.

Harris, J. D. (1943). Habituatory response decrement in the intact organism. Psychological Bulletin, 40(6):385.

Hassien, A. M., Shue, F., Bernier, B. E., and Drew, M. R. (2020). A mouse model of stress-enhanced fear learning demonstrates extinction-sensitive and extinction-resistant effects of footshock stress. Behavioural Brain Research, 379.

Hersman, S., Hoffman, A. N., Hodgins, L., Shieh, S., Lam, J., Parikh, A., and Fanselow, M. S. (2019). Cholinergic Signaling Alters Stress-Induced Sensitization of Hippocampal Contextual Learning. Frontiers in Neuroscience, 13.

Jász, A., Biró, L., Buday, Z., Király, B., Szalárdy, O., Horváth, K., Komlósi, G., Bódizs, R., Kovács, K. J., Diana, M. A., et al. (2025). Persistently increased post-stress activity of paraventricular thalamic neurons is essential for the emergence of stress-induced alterations in behaviour. PLoS biology, 23(1):e3002962.

Kamprath, K. and Wotjak, C. T. (2004). Nonassociative learning processes determine expression and extinction of conditioned fear in mice. Learning & Memory, 11(6):770–786.

Kim, E. J., Horovitz, O., Pellman, B. A., Tan, L. M., Li, Q., Richter-Levin, G., and Kim, J. J. (2013). Dorsal periaqueductal gray-amygdala pathway conveys both innate and learned fear responses in rats. Proceedings of the National Academy of Sciences, 110(36):14795–14800.

Kirouac, G. J. (2015). Placing the paraventricular nucleus of the thalamus within the brain circuits that control behavior. Neuroscience & Biobehavioral Reviews, 56:315–329.

Kolaj, M., Doroshenko, P., Cao, X. Y., Coderre, E., and Renaud, L. (2007). Orexin-induced modulation of statedependent intrinsic properties in thalamic paraventricular nucleus neurons attenuates action potential patterning and frequency. Neuroscience, 147(4):1066–1075.

Kooiker, C. L., Birnie, M. T., Floriou-Servou, A., Ding, Q., Thiagarajan, N., Hardy, M., and Baram, T. Z. (2024). Paraventricular thalamus neuronal ensembles encode early-life adversity and mediate the consequent sex-dependent disruptions of adult reward behaviors. bioRxiv.

Li, S. and Kirouac, G. J. (2012). Sources of inputs to the anterior and posterior aspects of the paraventricular nucleus of the thalamus. Brain Structure and Function, 217(2):257–273.

Li, Y., Dong, X., Li, S., and Kirouac, G. J. (2014). Lesions of the posterior paraventricular nucleus of the thalamus attenuate fear expression. Frontiers in behavioral neuroscience, 8:94.

Li, Y., Li, S., Wei, C., Wang, H., Sui, N., and Kirouac, G. J. (2010). Changes in emotional behavior produced by orexin microinjections in the paraventricular nucleus of the thalamus. Pharmacology Biochemistry and Behavior, 95(1):121–128.

Lissek, S. and Meurs, B. v. (2015). Learning models of PTSD: Theoretical accounts and psychobiological evidence. International Journal of Psychophysiology, 98(3):594–605.

Ma, J., Hoffmann, J. d., Kindel, M., Beas, B. S., Chudasama, Y., and Penzo, M. A. (2021). Divergent projections of the paraventricular nucleus of the thalamus mediate the selection of passive and active defensive behaviors. Nature Neuroscience, pages 1–12.

McFarlane, A. C. (2010). The long-term costs of traumatic stress: intertwined physical and psychological consequences. World Psychiatry, 9(1):3.

McGinty, J. F. and Otis, J. M. (2020). Heterogeneity in the Paraventricular Thalamus: The Traffic Light of Motivated Behaviors. Frontiers in Behavioral Neuroscience, 14:590528.

Nishimura, K. J., Poulos, A. M., Drew, M. R., and Rajbhandari, A. K. (2022). Know thy SEFL: Fear sensitization and its relevance to stressor-related disorders. Neuroscience and biobehavioral reviews, 142:104884.

Padilla-Coreano, N., Do-Monte, F. H., and Quirk, G. J. (2011). A time-dependent role of midline thalamic nuclei in the retrieval of fear memory. Neuropharmacology, 62(1):457–463.

Pennington, Z. T., Anderson, A. S., and Fanselow, M. S. (2017). The ventromedial prefrontal cortex in a model of traumatic stress: Fear inhibition or contextual processing? Learning and Memory, 24(9):400–406.

Pennington, Z. T., LaBanca, A. R., Sompolpong, P., Abdel-Raheim, S. D., Ko, B., Wick, Z. C., Feng, Y., Dong, Z., Francisco, T. R., Bacon, M. E., Chen, L., Fulton, S. L., Maze, I., Shuman, T., and Cai, D. J. (2024). Dissociable contributions of the amygdala and ventral hippocampus to stress-induced changes in defensive behavior. bioRxiv, page 2023.02.27.530077.

Penzo, M. A., Robert, V., Tucciarone, J., Bundel, D. D., Wang, M., Aelst, L. V., Darvas, M., Parada, L. F., Palmiter, R. D., He, M., Huang, Z. J., and Li, B. (2015). The paraventricular thalamus controls a central amygdala fear circuit. Nature, 519(7544):455–459.

Perusini, J. N., Meyer, E. M., Long, V. A., Rau, V., Nocera, N., Avershal, J., Maksymetz, J., Spigelman, I., and Fanselow, M. S. (2016). Induction and Expression of Fear Sensitization Caused by Acute Traumatic Stress. Neuropsychopharmacology, 41(1):45–57.

Pliota, P., Böhm, V., Grössl, F., Griessner, J., Valenti, O., Kraitsy, K., Kaczanowska, J., Pasieka, M., Lendl, T., Deussing, J. M., and Haubensak, W. (2018). Stress peptides sensitize fear circuitry to promote passive coping. Molecular Psychiatry, pages 1–14.

Quiñones-Laracuente, K., Vega-Medina, A., and Quirk, G. J. (2021). Time-Dependent Recruitment of Prelimbic Prefrontal Circuits for Retrieval of Fear Memory. Frontiers in Behavioral Neuroscience, 15:665116.

Rau, V., DeCola, J. P., and Fanselow, M. S. (2005). Stress-induced enhancement of fear learning: An animal model of posttraumatic stress disorder. Neuroscience & Biobehavioral Reviews, 29(8):1207–1223.

Sato, M., Ito, M., Nagase, M., Sugimura, Y. K., Takahashi, Y., Watabe, A. M., and Kato, F. (2015). The lateral parabrachial nucleus is actively involved in the acquisition of fear memory in mice. Molecular Brain, 8(1):22.

Siegmund, A. and Wotjak, C. T. (2007a). A mouse model of posttraumatic stress disorder that distinguishes between conditioned and sensitised fear. Journal of Psychiatric Research, 41(10):848–860.

Siegmund, A. and Wotjak, C. T. (2007b). Hyperarousal does not depend on trauma-related contextual memory in an animal model of Posttraumatic Stress Disorder. Physiology & Behavior, 90(1):103–107.

Sillivan, S. E., Joseph, N. F., Jamieson, S., King, M. L., Chévere-Torres, I., Fuentes, I., Shumyatsky, G. P., Brantley, A. F., Rumbaugh, G., and Miller, C. A. (2017). Susceptibility and Resilience to Posttraumatic Stress Disorder–like Behaviors in Inbred Mice. Biological Psychiatry, 82(12):924–933.

Stam, R. (2007a). PTSD and stress sensitisation: A tale of brain and body Part 1: Human studies. Neuroscience & Biobehavioral Reviews, 31(4):530–557.

Stam, R. (2007b). PTSD and stress sensitisation: A tale of brain and body Part 2: Animal models. Neuroscience & Biobehavioral Reviews, 31(4):558–584.

Temme, S. J., Bell, R. Z., Pahumi, R., and Murphy, G. G. (2014). Comparison of inbred mouse substrains reveals segregation of maladaptive fear phenotypes. Frontiers in behavioral neuroscience, 8:282.

Wang, T., Yan, R., Zhang, X., Wang, Z., Duan, H., Wang, Z., and Zhou, Q. (2023). Paraventricular thalamus dynamically modulates aversive memory via tuning prefrontal inhibitory circuitry. Journal of Neuroscience, 43(20):3630–3646.

Wang, Y., Pan, Y., Cai, Z., Lei, C., Guo, X., Cui, D., Yuan, Y., Lai, B., and Zheng, P. (2021). Inputs from paraventricular nucleus of thalamus and locus coeruleus contribute to the activation of central nucleus of amygdala during context-induced retrieval of morphine withdrawal memory. Experimental Neurology, 338:113600.

Wang, Z., Wang, Z., and Zhou, Q. (2024). Modulation of learning safety signals by acute stress: paraventricular thalamus and prefrontal inhibition. Neuropsychopharmacology, pages 1–13.

Zhu, Y., Nachtrab, G., Keyes, P. C., Allen, W. E., Luo, L., and Chen, X. (2018). Dynamic salience processing in paraventricular thalamus gates associative learning. Science, 362(6413):423–429.

Zhu, Y.-B., Wang, Y., Hua, X.-X., Xu, L., Liu, M.-Z., Zhang, R., Liu, P.-F., Li, J.-B., Zhang, L., and Mu, D. (2022). PBN-PVT projections modulate negative affective states in mice. eLife, 11:e68372.

